# *Enterococcus faecalis* AHG0090 is a genetically tractable bacterium and produces a secreted peptidic bioactive that suppresses NF-kB activation in human gut epithelial cells

**DOI:** 10.1101/275719

**Authors:** Páraic Ó Cuív, Rabina Giria, Emily C. Hoedt, Michael A. McGuckin, Jakob Begun, Mark Morrison

## Abstract

*Enterococcus faecalis* is an early coloniser of the human infant gut and contributes to the development of intestinal immunity. To better understand the functional capacity of *E. faecalis* we constructed a broad host range RP4 mobilisable vector, pEHR513112, that confers chloramphenicol resistance and used a metaparental mating approach to isolate *E. faecalis* AHG0090 from a faecal sample collected from a healthy human infant. We demonstrated that *E. faecalis* AHG0090 is genetically tractable and could be manipulated using traditional molecular microbiology approaches. *E. faecalis* AHG0090 was comparable to the gold-standard anti-inflammatory bacterium *Faecalibacterium prausnitzii* A2-165 in its ability to suppress cytokine mediated NF-КB activation in human gut derived LS174T goblet cell-like and Caco-2 enterocyte-like cell lines. *E. faecalis* AHG0090 and *F. prausnitzii* A2-165 produced secreted low molecular weight NF-КB suppressive peptidic bioactives. Both bioactives were sensitive to heat and proteinase K treatments although the *E. faecalis* AHG0090 bioactive was more resilient to both forms of treatment. As expected, *E. faecalis* AHG0090 suppressed IL-1β induced NF-КB-p65 subunit nuclear translocation and expression of the NF-КB regulated genes IL-6, IL-8 and CXCL-10. Finally, we determined that *E. faecalis* AHG0090 is distantly related to other commensal strains and likely encodes niche factors that support effective colonisation of the infant gut.

## Introduction

The human gut represents the largest mucosal surface area and is the largest immune organ of the body (Chassaing et al., 2014). Full-term infants are born with a competent but immature immune system that must respond appropriately to the inevitable exposure to microbes that occurs following birth. The infant microbiota is derived principally from the maternal microbiota (Dominguez-Bello et al., 2010) and the early colonisers of the gut play a critical role in priming mucosal immunity and establishing a homeostatic relationship with the host (Chung and Kasper, 2010;Fulde and Hornef, 2014).

*Enterococcus faecalis* is one of the most abundant colonisers of the infant gastrointestinal tract (Hopkins et al., 2005;Kirtzalidou et al., 2012) and together with other Enterococci, Staphylococci and Enterobacteria helps reduce the gut environment to facilitate subsequent colonisation by obligate anaerobes (Adlerberth, 2008;Wopereis et al., 2014). Select *E. faecalis* infant derived strains also possess immunomodulatory capacities (Wang et al., 2008) and exert anti-inflammatory activities by modulating the nuclear factors kappa B (NF-КB), mitogen activated protein kinase and peroxisome proliferator-activated receptor-γ1 regulated pathways (Are et al., 2008;Wang et al., 2014). Some of the immunomodulatory factors produced by *E. faecalis* have been identified (Brosnahan et al., 2013;Zou et al., 2014;Im et al., 2015) however the extent of immunomodulatory capacities amongst non-pathogenic gut strains, and the identity of the bioactives that underpin them, remains largely unknown.

Taken together, we hypothesised the infant gut microbiota would be a fertile source of immunomodulatory bioactive factors with potential prophylactic or therapeutic applications. We previously reported a method termed metaparental mating that enables the rapid and directed isolation of genetically tractable human gut bacteria (Ó Cuív et al., 2015). In this study, we describe the isolation of *E. faecalis* AHG0090 and demonstrate that similar to *Faecalibacterium prausnitzii* A2-165 (Quevrain et al., 2016;Breyner et al., 2017), it produces a potent peptidic bioactive that suppresses NF-КB activation. Finally, we demonstrate that *E. faecalis* AHG0090 can be manipulated using traditional molecular techniques providing new opportunities to dissect the functional capacity of the human gut microbiota.

## Materials & Methods

### Growth and culture conditions

The recipient cultures for metaparental mating were prepared by inoculating Brain Heart Infusion (BHI, Difco(tm)) supplemented with colistin sulfate with a raw stool sample collected from a healthy 2-year-old female child. The donor had not taken antibiotics during the 3-month period prior to collection. The child was recruited as part of a study into the link between the gut microbiota and type 1 diabetes susceptibility. All study samples were collected in accordance with the recommendations of the Mater Health Services Human Research Ethics Committee (HREC/13/MHS/21/AM02). All subjects gave written informed consent in accordance with the Declaration of Helsinki, with written consent provided from parents or legal guardians for all subjects <13 years. The protocol was approved by the Mater Health Services Human Research Ethics Committee. *E. faecalis* was cultured in BHI and the *Escherichia coli* ST18 donor strain for metaparental mating was cultured in BHI supplemented with δ-aminolevulinic acid (100 μg.ml-1). The *E. coli* cloning strains were grown in LB and *F. prausnitzii* A2-165 was cultured in anaerobic Reinforced Clostridial Medium (RCM, Oxoid(tm)) buffered with salt solutions 2 and 3 (McSweeney et al., 2005). *F. prausnitzii* cultures were routinely manipulated in a Coy vinyl anaerobic chamber with an oxygen free atmosphere (85% N2:10% CO2:5% H2). Both *E. coli* ST18 and JM109 competent cells were prepared by the rubidium chloride method (Hanahan, 1985) while InvitrogenTM E. coli INVα F′ competent cells were purchased from ThermoFisher Scientific. The bacterial growth media were supplemented with erythromycin (100 μg.ml-1), chloramphenicol (20 μg.ml-1) or colisitin sulfate (20 μg.ml-^1^) as appropriate.

Bacterial growth was measured by the increase in optical density at 600 nm (OD600nm). Specific growth rates (μ [hours-^1^]) were calculated by log^10^ transformation of the OD600nm measurements and plotting a trendline (R^2^>0.97) for the linear phase of growth corresponding to exponential growth phase. Then μ was calculated using the equation: μ = (slope of the line * 2.3).

### Vector construction and manipulation

To construct pEHR513111, *catP* was PCR amplified from ^pJIR1456^ (Lyras and Rood, 1998) with primers (Pf5’ GAT CGT TTA AAC AGT GGG CAA GTT GAA AAA TTC AC; Pr5’ GAT CCC TGC AGG TTA GGG TAA CAA AAA ACA CCG TAT TTC TAC) that introduced unique PmeI and SbfI restriction sites. The digested *catP* was then cloned into pEHR512111, replacing *erm*, and generating pEHR513111. The pEHR513112 vector was subsequently constructed by cloning *cphy_3290-evolglow-C-Bs2* from pEHR512112 (Ó Cuív et al., 2015) into the multiple cloning site of pEHR513111 as an EcoRI-HindIII fragment. The pEHR513111 and pEHR513112 vectors were confirmed by restriction digest analysis and the sequences compiled using publicly available sequences. When appropriate, the pEHR513112 vector was cured from *E. faecalis* AHG0090 by overnight growth in BHI medium supplemented with acridine orange (1-8 μg.ml-^1^) and then plating on BHI medium. Single colonies were replica plated onto BHI medium with or without chloramphenicol to identify naïve *E. faecalis* AHG0090 isolates.

Plasmid DNA was extracted from *E. faecalis* cultures using a modified alkaline lysis method (Green and Sambrook, 2016). Briefly, 1 ml of *E. faecalis* culture was centrifuged and the cells were resuspended in 200 μl of Solution 1. The cell suspension was supplemented with 1 μl of mutanolysin (20 U/μl) and 10 μl lysozyme (200 mg/ml) and incubated at 37°C for up to 1 hour. Next, 200 μl of Solution 2 was added, and the mixture was incubated on ice for 5 min. Then, 200 μl of Solution 3 was added, and the mixture was incubated on ice for 10 min. The cell debris were removed by centrifugation and the clarified cell lysate was recovered and extracted with phenol:chloroform:isoamyalcohol (25:24:1). The aqueous phase was transferred to a fresh tube and plasmid DNA was precipitated using isopropanol, washed with 70% (v/v) ethanol and then resuspended in TE buffer. Plasmid DNA was similarly prepared from *E. coli* except that the treatments with mutanolysin and lysozyme, and the incubations on ice were not performed.

### Mating procedures

The *E. coli* ST18 donor strain for metaparental and bi-parental matings was grown in BHI medium supplemented with δ-aminolevulinic acid and chloramphenicol. The metaparental mating experiments were performed essentially as previously described (Ó Cuív et al., 2015) with two exceptions. First, the mating mix and controls were spotted directly onto the surface of BHI agar rather than plating on a nylon filter. Second, 100 μl of mating mix and controls were transferred to fresh BHI broth and grown for 5 hours before selecting for transconjugants. Biparental matings were similarly performed except that the 5 hour outgrowth was not done. Transconjugants were recovered on BHI medium supplemented with chloramphenicol and colisin sulfate.

### Microscopy

Naïve and transconjugants strains of *E. faecalis* AHG0090 were examined with an Olympus BX 63 microscope fitted with an Xcite LED light source and fluorescence filter cube U-FBN (excitation 470–495 nm, emission 510 nm). Images were captured using a DP80 camera and the Olympus cellSens modular imaging software platform and ImageJ software package (http://imagej.nih.gov/ij/) were used for visualisation and processing.

### *E. faecalis rrs* sequencing

The *rrs* gene was PCR amplified using the primers 27F and 1492R (Lane, 1990) as previously described (Ó Cuív et al., 2011) and sequenced at the Australian Genomic Research Facility (Brisbane, Australia) using primers 530F and 907R (Lane, 1990). The individual sequence reads were trimmed to remove low quality bases and assembled using deFUME (van der Helm et al., 2015). The assembled sequence was then aligned against the Ribosomal Database Project (Cole et al., 2014) core set of aligned *rrs* sequences and *E. faecalis* AHG0090 was identified using the Classifier and SeqMatch functions.

### Genome sequencing and analysis

High molecular weight DNA was prepared from the *E. faecalis* AHG0090 metaparental mating isolate as previously described (Ó Cuív et al., 2011). The DNA was then quantified using the QuantiFluor ONE dsDNA system according to manufacturer’s instructions (Promega, Australia) and the integrity of the DNA was determined by agarose gel electrophoresis. The genome was shotgun sequenced using the Illumina NextSeq 500 system (2 x 150bp High Output kit) with v2 chemistry. The sequence data were quality checked, filtered and then *de novo* assembled using the SPAdes assembler v 3.10.1 (Bankevich et al., 2012). Genome sequencing quality was evaluated with CheckM, which estimates the input files for completeness and contamination based on the phylogenetic assignment of a broad set of marker genes (Parks et al., 2015). The *E. faecalis* AHG0090 contigs were ordered using Mauve (Darling et al., 2010) with the closed *E. faecalis* V583 genome sequence used as a reference. The Mauve generated assembly was submitted to the RAST annotation pipeline and the results were examined in the SEED Viewer (Aziz et al., 2008). Genome based phylogeny was obtained using GTDB (https://github.com/Ecogenomics/GTDBLite), built from the concatenation of 120 universal bacterial-specific marker genes (Parks et al., 2017). Tree inference was performed with FastTree v2.1.7 (Price et al., 2010) and included all genomes in IMG v4.510 (Markowitz et al., 2012). The resulting tree was imaged using ARB v6.0.6 (Ludwig et al., 2004). To identify candidate plasmids, the fastq files were mapped to the *E. faecalis* AHG0090 genome assembly using BamM v1.7.3 (http://ecogenomics.github.io/BamM/) to determine the coverage profiles for each contig. The average coverage was then calculated and contigs with >1000x coverage were identified as candidate plasmids. Additionally, we used PlasmidSPAdes to assemble plasmids from whole genome sequencing data (Antipov et al., 2016). The candidate plasmids were manually curated to determine if they could be closed and compared to other plasmids using Blastn. This Whole Genome Shotgun project was deposited at DDBJ/EMBL/GenBank under the accession PDUN00000000. The version described in this paper is the first version, PDUN01000000.

### Measurement of *E. faecalis* immunomodulatory activities

The immunomodulatory potential of *E. faecalis* was assessed using LS174T-NF-КB*luc* goblet cell like and Caco-2-NF-КB*luc* enterocyte like reporter cell lines (Ó Cuív et al., 2017). Briefly, three individual *E. faecalis* AHG0090 colonies were established as independent cultures with BHI broth. Following overnight growth, each individual culture was used to initiate duplicate 50 ml BHI broth cultures at a starting OD600nm of 0.01 (n=6 cultures, consisting of n=3 independent biological replicates with n=2 technical replicates each). The OD600nm of the cultures was monitored longitudinally and 5 ml of cultures was harvested from each broth culture at early exponential, mid-exponential, early stationary and late stationary phase of growth. At each collection, 1.5 ml of each culture was centrifuged at 16,000 x g for 5 minutes and 0.5 ml of the cell-free supernatant fraction was transferred to fresh tubes and stored at −30oC as single-use aliquots.

For the immunomodulatory assays, 96-well microtiter plates were seeded with 20,000 LS174T-NF-kB*luc* or Caco-2-NF-kB*luc* reporter cells per well as previously described (Ó Cuív et al., 2017). The ability of the cell free bacterial supernatants to suppress NF-КB activation in LS174T-NF-kB*luc* was assessed by adding supernatant (10% v/v in complete DMEM medium) to the cells and incubating for 30 minutes at 37°C. The LS174T and Caco-2 reporter cell lines were then stimulated with TNFα (25ng.ml-1) or IL-1β (50 ng.ml-1) respectively in the presence of 10% v/v supernatant for 4 hours before assessing luciferase activity. The ability of the supernatants to suppress activation was compared to the NF-КB inhibitor indole-3-carbinol (I3C, 5 μM). NF-κB driven luciferase expression was assessed using the PierceTM Firefly Luc One-Step Glow Assay Kit (ThermoFisher Scientific) according to the manufacturer’s instructions. The cytotoxicity of the supernatants was assessed using Cell Proliferation Reagent WST-1 (Sigma Aldirch) according to the manufacturer’s instructions.

### Nuclear translocation immunofluorescence assays

Glass coverslips in a 12 well-plate were seeded with 20,000 Caco-2 cells per well and cultured overnight. Cell-free culture supernatants harvested from mid-exponential phase cultures were added (10% v/v) to the Caco-2 cells and incubated for 30 min, and then stimulated with IL-1β (25ng.ml-1) for 1 hour. Cells were also treated with BHI medium and I3C alone. The cells were then processed and analysed as previously described (Ó Cuív et al., 2017).

### Quantitative reverse transcriptase PCR (qRT-PCR) assays

A 12 well-plate was seeded with 50,000 Caco-2 cells per well and cultured overnight. Cell-free culture supernatants harvested from mid-exponential phase cultures were added (10% v/v) to the Caco-2 cells and incubated for 30 min, and then stimulated with IL-1β (25ng.ml-^1^) for 6 hours. Cells were also treated with I3C and BHI medium alone. Total RNA was isolated and the expression of the NF-КB dependent genes IL-6, IL-8 and CXCL10 was assessed as previously described (Ó Cuív et al., 2017), except that different primers were used for IL-6 (Pf5’ CCA CTC ACC TCT TCA GAA CG; Pr5’ CAT CTT TGG AAG GTT CAG GTT G) (Noss et al., 2015).

## Results

### Isolation of *E. faecalis* AHG0090

To recover genetically tractable facultative anaerobic *Firmicutes* bacteria from the healthy infant human gut we produced a microbial enrichment culture from human stool and determined that the addition of chloramphenicol completely inhibited growth of candidate recipients on BHI medium under aerobic conditions. Consequently, we constructed a vector, pEHR513111, carrying *catP* and this vector was further modified by cloning the *evoglow-C-Bs2* under the control of the *Clostridium phyofermentans* ISDg *cphy_3290* promoter generating pEHR513112 (Figure 1A).

**Figure 1.**
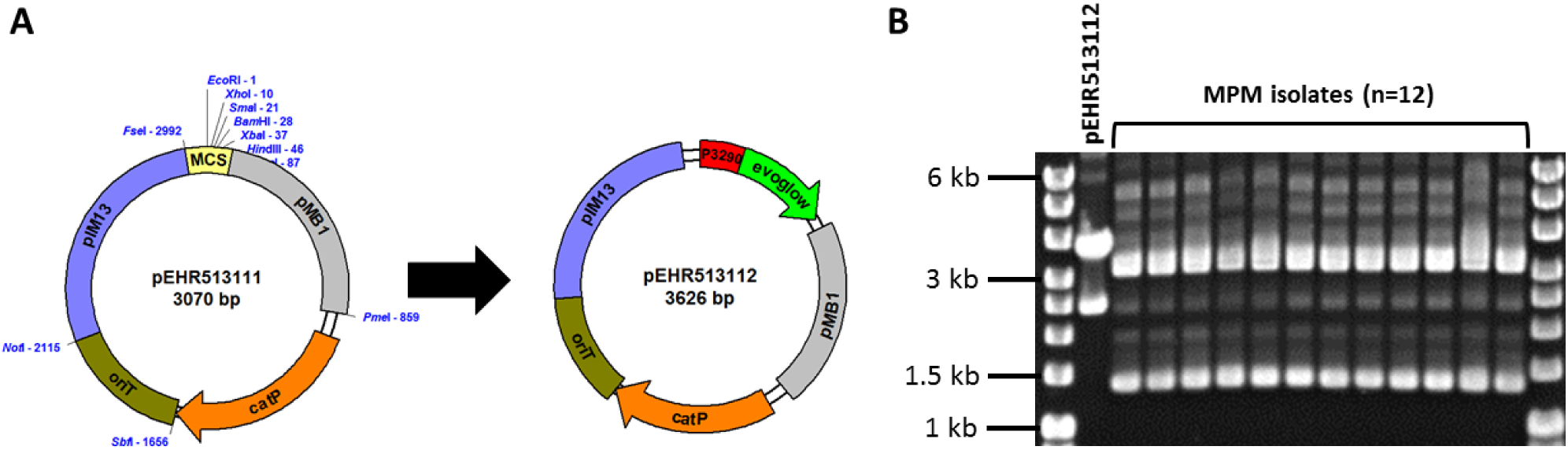
The pEHR513111 plasmid carrying *catP* confers chloramphenicol resistance on its host. The pEHR513112 plasmid carries *evoglow-C-Bs2* under the control of the *C. phyofermentans* ISDg *cphy_3290* promoter. **B.** Plasmid profiling of the transconjugants recovered by metaparental mating (MPM isolates). The pEHR513112 plasmid carried by the transconjugants was identified by comparison with plasmid DNA prepared from *E. coli* (pEHR513112).

Following metaparental mating we recovered 14 transconjugants and assessed the clonality of 12 isolates by plasmid profiling. All 12 isolates carried a plasmid of comparable molecular weight to pEHR513112 in addition to at least two other plasmids (∼1.4 kb closed covalent circular form, ∼3.2 kb closed covalent circular form) (Figure 1B). However, the plasmid DNA profiles of all 12 isolates were virtually identical suggesting they were clonal. Based on these observations we chose one transconjugant for further analysis and produced 1473 bp of 16S rRNA sequence. Based on this sequence we determined the isolate was affiliated with the *Enterococcus faecalis* taxon, and hereafter it is referred to as *E. faecalis* AHG0090.

### *E. faecalis* AHG0090 is genetically tractable

We examined whether *E. faecalis* AHG0090 can be genetically manipulated using traditional techniques in molecular microbiology. *E. faecalis* AHG0090 was grown in BHI broth supplemented with acridine orange to cure pEHR513112. The addition of acridine orange up to 8 μg.ml-^1^ did not affect growth however all the colonies recovered on BHI medium were sensitive to chloramphenicol suggesting they had lost pEHR513112. Plasmid DNA prepared from *E. faecalis* AHG0090 recovered by metaparental mating carried a plasmid with the same molecular weight as pEHR513112 however, this plasmid was absent from naïve *E. faecalis* AHG0090 (Figure 2A). We next confirmed that naïve *E. faecalis* AHG0090 was genetically tractable by using it as the recipient in a biparental mating with *E. coli* ST18 carrying pEHR513112. Using our standard biparental mating protocol we recovered transconjugants and achieved a conjugation efficiency of 3.83 x 10-^7^ transconjugants per recipient. As expected, the biparental mating derived transconjugant carried the pEHR513112 plasmid band (Figure 2A) and plasmid recovery experiments from the metaparental mating and re-transformed isolates confirmed the plasmids were stably maintained (Figure 2B). Consistent with these observations the re-transformed but not naïve strain was fluorescent (Figure 2C). Notably, the endogenous plasmids were unaffected by the acridine orange treatment suggesting that they are stably maintained.

**Figure 2.**
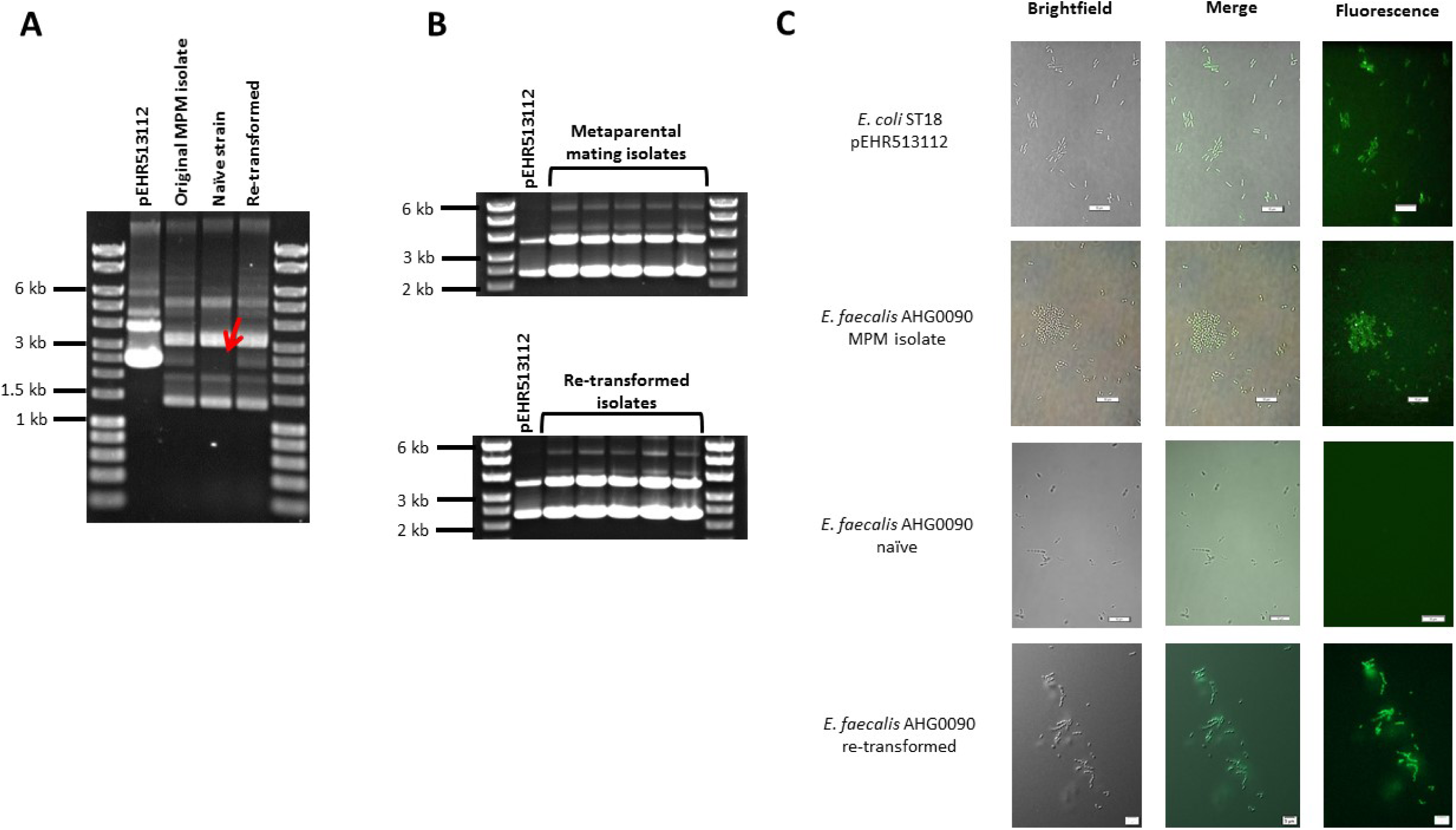
*E. faecalis* AHG0090 is genetically tractable and can be manipulated using standard molecular microbiology approaches. The pEHR513112 plasmid was readily identifiable in *E. faecalis* AHG0090 recovered by metaparental mating (original MPM isolate) and following the re-introduction of the plasmid by biparental mating (re-transformed), but was lost following treatment with acridine orange (naïve strain). The absence of pEHR513112 in the naïve strain is indicated by a red arrow. **B.** The pEHR513112 plasmid is stably maintained in *E. faecalis* AHG0090. pEHR513112 plasmids recovered from *E. faecalis* AHG0090 transconjugants produced by metaparental mating and following re-transformation of the naïve strain were examined by agarose gel electrophoresis to identify any major structural deletions and re-arrangements. **C.** *E. faecalis* strains carrying pEHR513112 are fluorescent. *E. coli* ST18 carrying pEHR513112 and naïve and transconjugant *E. faecalis* AHG0090 strains were analysed using brightfield and fluorescence microscopy. The scale bars represent 10?μm.

### *E. faecalis* AHG0090 produces an NF-КB suppressive peptidic bioactive

Given the immunomodulatory activity previously ascribed to *E. faecalis* isolates we examined the ability of *E. faecalis* AHG0090 to suppress cytokine mediated epithelial NF-КB activation using our LS174T and Caco-2 reporter cell lines. *E. faecalis* AHG0090 was grown in BHI medium and achieved a specific growth rate of 1.63±0.14 h-1 (Growth rate±SD) during exponential growth phase and a maximum recorded yield of 3.74±0.07 (OD600 ±SD) following 8 hours of growth (Figure 3A). Cell free supernatants were harvested from early exponential, mid-exponential, early stationary and late stationary phase cultures as the closely related bacterium *Lactobacillus plantarum* produces immunomodulins that inhibit IFNγ production in a growth phase dependent manner (Zvanych et al., 2014). Culture supernatant harvested from all four-time points suppressed NF-КB activation in both cell lines although the extent of suppression was greatest with supernatants harvested from mid-exponential phase onward (Figure 3B).

**Figure 3.**
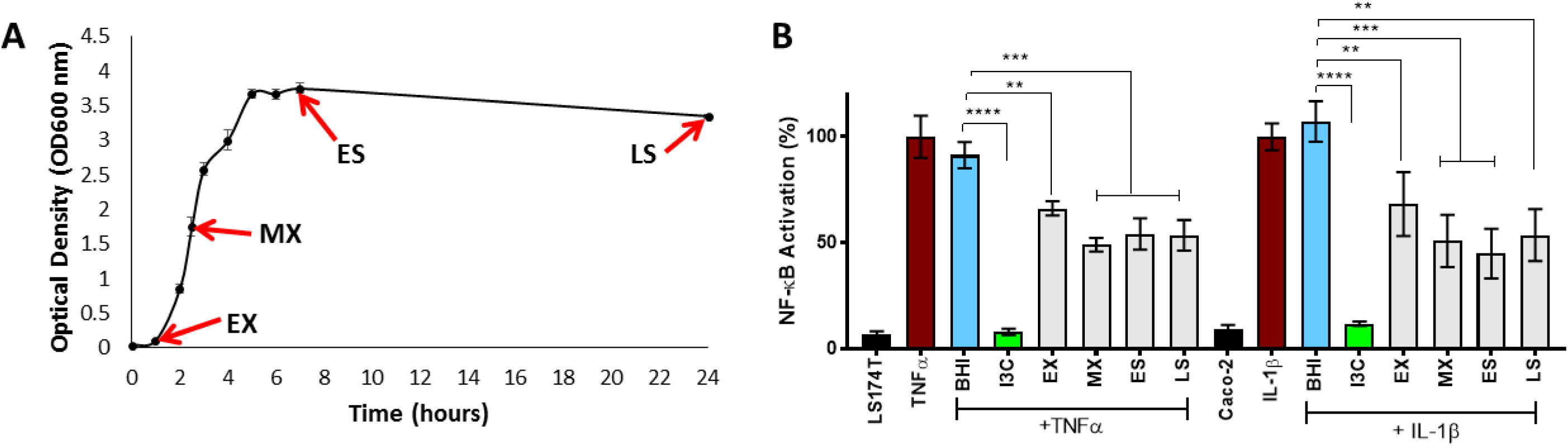
Harvesting of *E. faecalis* AHG0090 cell-free culture supernatants. *E. faecalis* AHG0090 was cultured in BHI medium and culture supernatants were harvested in early exponential (EX), mid-exponential (MX), early stationary (ES) and late stationary (LS) phase. Growth was recorded as the change in optical density over time (hours). **B.** Characterisation of the NF-КB suppressive capacity of *E. faecalis* AHG0090. The effects of the *E. faecalis* AHG0090 culture supernatants on NF-КB activation in the LS174T-NF-КB*luc* and Caco-2-NF-КB*luc* reporter cell lines were measured by the luciferase assay. The extent of NF-КB activation was assessed after 6 hours stimulation with IL-1β and baseline suppression of the reporter gene was assessed using sterile BHI medium (** *p*<0.01, *** *p*<0.001, **** *p*<0.0001 as determined by one-way ANOVA with Dunnett’s multiple comparison test).

We compared the NF-КB suppressive capacity of *E. faecalis* AHG0090 to the model anti-inflammatory gut bacterium *F. prausnitzii* A2-165 and determined that both strains suppressed NF-КB activation to a similar extent in the LS174T and Caco-2 reporter cell lines (Figure 4A). Critically, the *E. faecalis* AHG0090 and *F. prausnitzii* A2-165 cell free supernatants did not exert cytotoxic effects. The *E. faecalis* AHG0090 culture supernatant was fractionated by passing it through a 3 kDa molecular weight cut-off filter and the NF-КB suppressive activity of the flow-through but not the retentate was similar to that of the unfractionated culture supernatant (Figure 4A, *p*>0.05). We next assessed the impact of heat and proteinase K treatments on the *F. prausnitzii* A2-165 and *E. faecalis* AHG0090 NF-КB suppressive bioactives. The activity of the *F. prausnitzii* A2-165 <3 kDa culture supernatant fraction was not significantly different to the RCM control following heat (57 and 97°C) or proteinase K treatment (Figure 4B, *p*>0.05), consistent with the NF-КB suppressive capacity of this bacterium being mediated by Mam derived peptides (Quevrain et al., 2016). The *E. faecalis* AHG0090 <3 kDa culture supernatant fraction displayed similar characteristics but still retained activity following treatment at 57°C for 30 min or proteinase K digestion for 1 hour when compared to the BHI control (Figure 4B, *p*<0.05). This activity was lost at higher temperatures or following longer heat treatment, and following extended proteinase K treatment (Figure 4B, *p*>0.05). Taken together, these data suggest *E. faecalis* AHG0090 secretes a low molecular weight NF-КB suppressive peptidic bioactive with differing properties to the *F. prausnitzii* Mam peptides.

**Figure 4.**
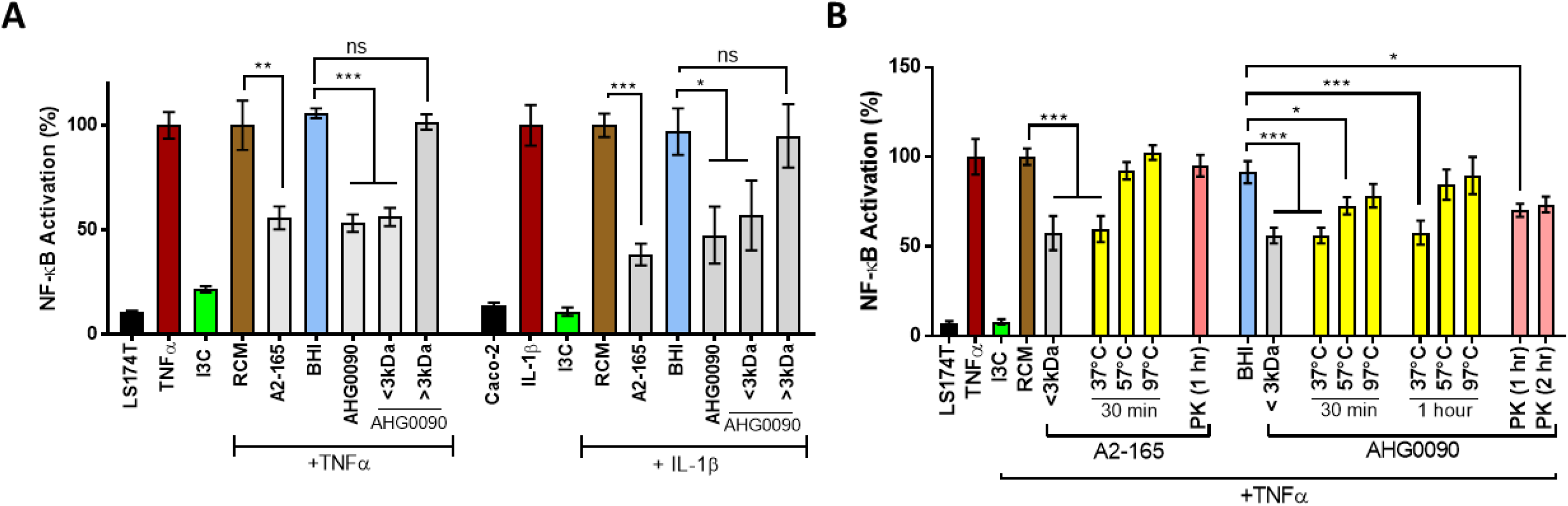
*F. prausnitzii* A2-165 and *E. faecalis* AHG0090 produce low molecular weight NF-КB suppressive bioactives. The extent of NF-КB activation was assessed after 6 hours stimulation of the LS174T-NF-kB*luc* and Caco-2-NF-kB*luc* reporter cell lines with TNFα and IL-1β respectively. Baseline suppression of the reporter gene was assessed using sterile RCM or BHI medium. The suppressive effects of the supernatants were assessed against the appropriate medium control. **B.** *F. prausnitzii* A2-165 and *E. faecalis* AHG0090 produce heat and proteinase K labile bioactives. The effect of the heat and proteinase K treatments was assessed using the LS174T-NF-КB*luc* reporter cell line. The suppressive effects of the supernatants were assessed against the appropriate medium control and significant differences are indicated. ns not significant, * *p*<0.05, ** *p*<0.01, *** *p*<0.001, **** *p*<0.0001 as determined by one-way ANOVA with Dunnett’s multiple comparison test.

### *E. faecalis* AHG0090 inhibits NF-КB-p65 subunit nuclear translocation and cytokine expression

Cytokine mediate activation of the NF-КB pathway results in nuclear translocation of the NF-КB-p65 subunit. We examined the ability of culture supernatant harvested from mid-exponential phase cultures of *E. faecalis* AHG0090 to suppress NF-КB–p65 subunit nuclear translocation. NF-КB–p65 subunit nuclear translocation induced by IL-1β in Caco-2 cells was unaffected by treatment with BHI medium. In contrast, *E. faecalis* AHG0090 mid-exponential phase culture supernatant treatment significantly reduced nuclear translocation in a similar fashion to the pharmacological inhibitor I3C (Figure 5A). As expected, I3C and *E. faecalis* AHG0090 culture supernatant suppressed expression of the NF-КB dependent genes IL-6 (*p*<0.05), IL-8 (*p*<0.01) and CXCL10 (*p*<0.001), as determined by qRT-PCR (Figure 5B).

### *E. faecalis* AHG0090 is adapted for gut colonisation

We sequenced the *E. faecalis* AHG0090 genome to provide insights into the factors supporting colonisation and persistence in the infant gut. We produced 2,925,542 bp of DNA sequence at 107x coverage. The sequenced data was assembled into 116 contigs providing a contig N50 of 144,336 bp and L50 of 8. Critically, the genome was assessed by CheckM as being essentially complete (99.63%) and free from contamination. The genome has a G+C content of 37.3% and is predicted to contain 2,929 protein-coding genes and 60 structural RNAs. Analysis of *E. faecalis* phylogeny using the GTDB revealed that *E. faecalis* AHG0090 clusters closely with three strains termed *E. faecalis* TX0630, TX0635 and TX0645, and distally from the *E. faecalis* type strains (*E. faecalis* ATCC19433 and 29200), and other gut commensal (e.g. *E. faecalis* PC1.1, 62 and Symbioflor1) and pathogenic (*E. faecalis* V583) strains (Figure 6A). Although *E. faecalis* is characterised by extensive horizontal gene transfer there is a high degree of synteny between *E. faecalis* AHG0090 and the closed commensal (*E. faecalis* 62) and pathogenic (*E. faecalis* V583) strains (Figure 6B).

We identified several plasmids in the *E. faecalis* AHG0090 genome sequence. pAHG0090c is a predicted to be 76,529 bp and is comprised of 11 contigs. It is predicted to encode 80 genes and displays sequence similarity and synteny to another plasmid from *E. faecalis* NKH15 (pMG2200 [106,527 bp], 99% identity and 72% coverage). Enterococcal plasmids are widely shared through horizontal transfer and we identified an aggregation substance encoding regulon that mediates efficient contact between donor and recipient bacteria to facilitate plasmid transfer (Bhatty et al., 2015), and adhesion to host cells (Olmsted et al., 1994;Vanek et al., 1999). We also identified two closed endogenous plasmid sequences in the genome sequence data. pAHG0090b is a 5,121 bp plasmid that exhibits sequence similarity to plasmids from *E. faecalis* 62 (EF62pA [5,143 bp], 99% identity and 100% coverage) (Brede et al., 2011) and *E. faecalis* S-86 (pS86 [5,149 bp], 99% identity and 99% coverage) (Martinez-Bueno et al., 2000). pAHG0090b has a G+C content of 37% and is predicted to encode 6 proteins. pAHG0090a is a 1,925 bp plasmid that exhibits extensive sequence similarity to a cryptic plasmid of similar size from *Enterococcus faecium* 226 (pMBB1 [1,932 bp], 96% identity and 67% coverage) (Wyckoff et al., 1996), and larger cryptic plasmids from *Lactococcus fermentum* KC5b (pKC5b [4,392 bp], 99% identity and 75% coverge) (Pavlova et al., 2002) and *Lactococcus lactis* (pCRL291.1 [4,640 bp], 89% identity and 70% coverage). pAHG0090a has a G+C content of 33% and is predicted to encode a single protein predicted to function in plasmid replication.

**Figure 5.**
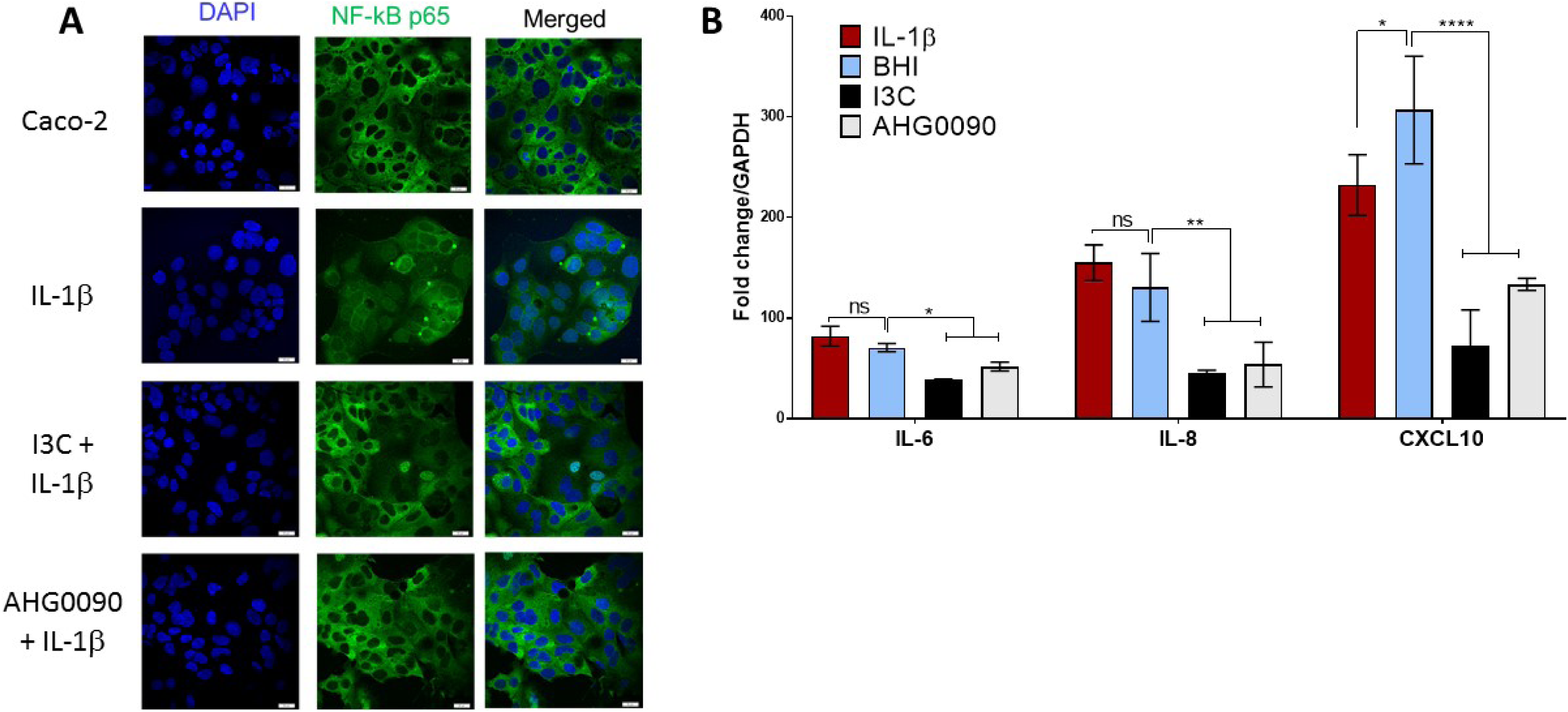
*E. faecalis* AHG0090 mid-exponential phase culture supernatant suppresses NF-КB-p65 subunit nuclear translocation. The cell nuclei and NF-КB-p65 subunit are shown in blue and green respectively. The nuclei for the BHI treated Caco-2 cells in the central panel are largely black revealing that treatment does not result in nuclear translocation. As expected, BHI treatment does not prevent NF-КB-p65 nuclear translocation following IL-1β treatment as indicated by green staining of the nuclei. In contrast, treatment with I3C and *E. faecalis* AHG0090 mid-exponential phase culture supernatant suppressed IL-1β induced NF-КB-p65 nuclear translocation. The scale bars represent 10 μm. **B.** *E. faecalis* AHG0090 mid-exponential phase culture supernatants suppress expression of NF-КB-p65 dependent cytokines. The expression of IL-6, IL-8 and CXCL10 was assessed by qRT-PCR. The data are normalized to GAPDH gene expression and presented as the fold-change relative to unstimulated cells. *E. faecalis* AHG0090 mid-exponential phase culture supernatants suppress expression of IL-6 (\**p*<0.05), IL-8 (\*\**p*<0.01) and CXCL10 (\*\*\*\**p*<0.0001) as determined using one-way ANOVA with Dunnett’s multiple comparison test.

**Figure 6.**
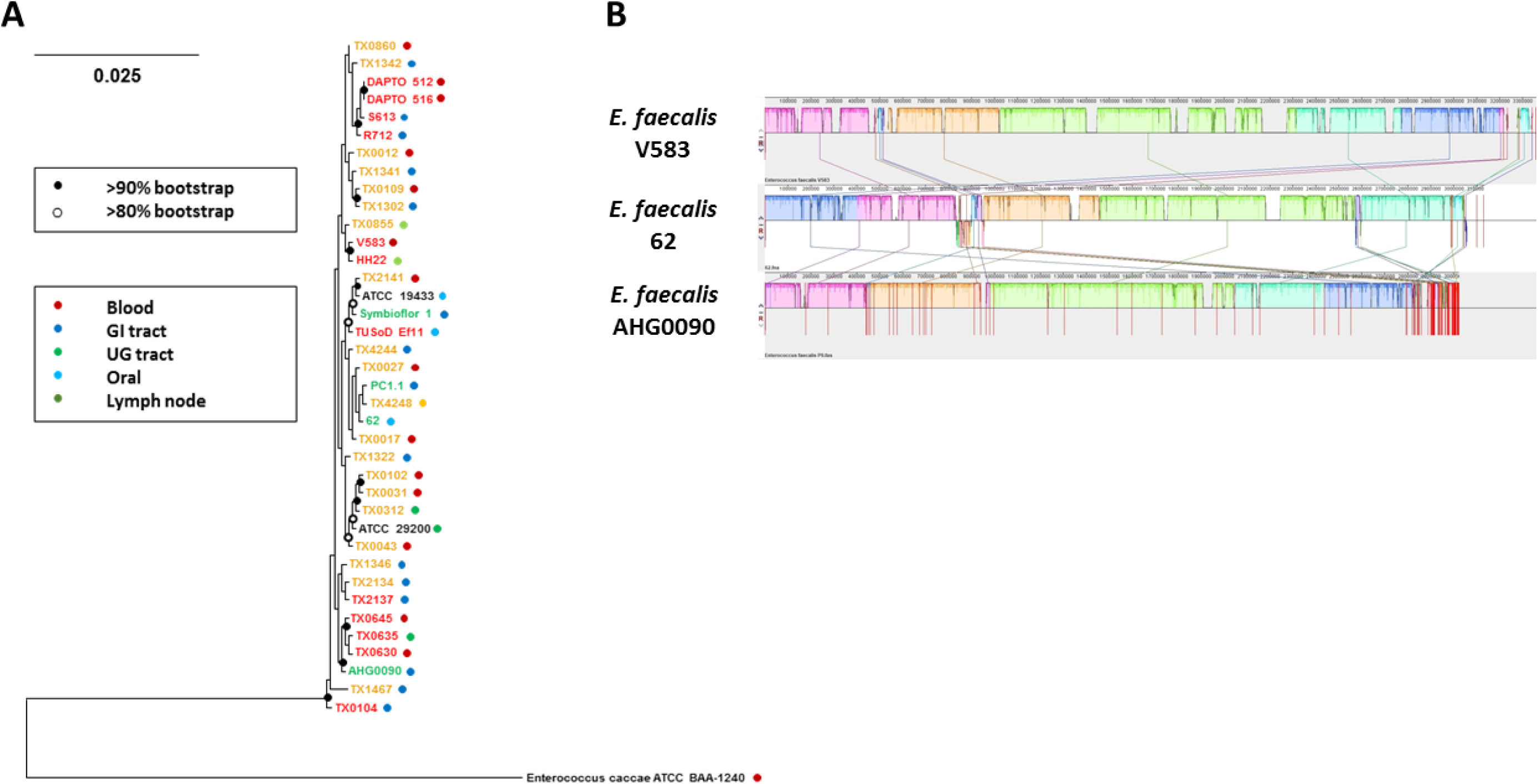
GTDB based phylogeny of *E. faecalis* as determined from the concatenation of 120 universal bacterial-specific marker genes. The source of individual commensal (green), uncharacterised (orange) and pathogenic (red) strains is indicated. The bootstrap values are indicated using a cut-off of >80 or >90%. **B.** The extent of genome synteny between *E. faecalis* AHG0090 and a representative pathogenic strain (*E. faecalis* V583) and commensal (*E. faecalis* 62) that have closed genome sequences. The red lines for *E. faecalis* V583 and 62 indicated boundaries of the chromosomes and plasmids whereas the red lines for *E. faecalis* AHG0090 indicate individual contig sequences.

We also identified pEHR513112 in the genome sequence. There were 4 differences between the compiled pEHR513112 sequence and the vector carried by *E. faecalis* AHG0090. We identified a G86T transversion in the *cphy_3290* promoter and a C2407T transition in *oriT* vector modules. We also identified a 1 bp deletion at the 5’ (AscI) end and a separate 11 bp deletion at the 3’ (PmeI) end of the pMB1 module. Notably, these deletions occurred within the primer sequences used to PCR amplify the pMB1 module.

The *E. faecalis* AHG0090 chromosome is predicted to encode a range of niche factors that likely support colonisation and persistence in the infant gut. Many Enterococci are non-motile and the ability to adhere to the host epithelium likely plays a role in preventing wash-out. We identified an Ebp-like pilus, the microbial surface component recognizing adhesive matrix molecules protein Ace and several adhesins including EfbA that mediate adhesion to host structural factors (e.g. collagen, fibrinogen and laminin) and support biofilm formation (Nallapareddy et al., 2000;Montealegre et al., 2015;Singh et al., 2015). *E. faecalis* AHG0090 also encodes several proteins that enable foraging of host glycans (Bohle et al., 2011;Garbe et al., 2014) and it also encodes both the GelE and SprE proteases that have been proposed to support the nutrient requirements of the bacterium by digesting host proteins and cells (Fisher and Phillips, 2009). We also identified several proteins that likely modulate interactions with the host immune system including an internalin like protein that may support intracellular persistence (Brinster et al., 2007) and a capsule that may contribute to immune evasion (Thurlow et al., 2009). Notably, we also identified a TIR domain protein previously shown to suppress MyD88 signalling and NF-КB activation by *E. faecalis* V583 (Zou et al., 2014). As expected, *E. faecalis* AHG0090 does not encode Mam like sequences and we did not identify any candidate genes and/or regulons likely to encode a low molecular weight NF-КB suppressive peptidic bioactive (e.g. bacteriocin CBT-SL5((Lee et al., 2008), the *E. faecalis* SL-5 bacteriocin CBT-SL5 is likely same as bacteriocin ESL5 which is produced by the same strain (Kang et al., 2009)) such as the bioactive we describe.

## Discussion

The early microbial colonisers of the gut help establish a homeostatic relationship between the host and its microbiota (Battersby and Gibbons, 2013;Fulde and Hornef, 2014). *E. faecalis* comprises part of the vaginal (Brosnahan et al., 2013;Nami et al., 2014) and breastmilk (Jimenez et al., 2008;Albesharat et al., 2011;Kozak et al., 2015) microbiota, and is widely shared between mothers and their infants. It is increasingly recognised that early life events (e.g. method of birth, feeding) modify risks for several chronic diseases (Renz-Polster et al., 2005;Ng et al., 2015) and this may be related at least in part to early differences in gut colonisation and immune modulation. Much remains to be discovered about the bacteria and bioactive factors that underpin these events, and whether they could be exploited to optimise health and appropriate establishment of gut mucosal immunity.

In this study, we describe the isolation of a genetically tractable *E. faecalis* strain from infant stool and demonstrate that it produces a potent NF-КB suppressive bioactive. The suppressive activity of *E. faecalis* AHG0090 was clearly apparent in early exponential phase culture supernatants and did not increase significantly from mid-exponential phase onward. This suggests the bioactive is produced in early growth and persists in the culture supernatant through the proceeding phases of growth. The closely related bacterium *Lactobacillus plantarum* WCFS1 also exerts NF-КB and IFNγ suppressive effects and produces bioactives in a growth phase dependent manner (van Baarlen et al., 2009;Zvanych et al., 2014), possibly as a response of the bacterium to increased nutrient limitation during the transition from mid-log to stationary phase. We determined that the ability of *E. faecalis* AHG0090 to suppress NF-КB in our reporter cell lines was comparable to that of *F. prausnitzii* A2-165. *F. prausnitzii* is widely regarded as a model anti-inflammatory fastidious gut bacterium and produces a 15 kDa protein termed Mam that underpins NF-КB suppression (Quevrain et al., 2016;Breyner et al., 2017). Mam derived peptides are detectable in *F. prausnitzii* culture supernatants however it has not yet been reported whether the NF-КB suppressive activity is mediated by Mam and/or its peptide derivatives. While this remains to be further explored, we showed that NF-КB activation was suppressed by the <3 kDa culture supernatant fraction, and the suppressive effect was abrogated by heat or proteinase K treatment. Taken together, this suggests suppression of NF-КB by *F. prausnitzii* in our assays was mediated by <3 kDa Mam derived peptides. Our data also suggests that the NF-КB suppressive activity of *E. faecalis* AHG0090 is mediated by a low molecular weight peptidic bioactive although it is more resilient to heat and proteinase K treatment than Mam derived peptides.

The *E. faecalis* AHG0090 genome sequence allowed us to readily predict “known” functionalities. Both commensal and pathogenic strains of *E. faecalis* have previously been shown to produce NF-КB suppressive factors (Brosnahan et al., 2013;Zou et al., 2014). *E. faecalis* V583 encodes a TIR domain containing protein, TcpF, that suppresses NF-КB by interfering with MyD88 signalling (Zou et al., 2014). NF-КB suppression by TcpF is dependent on contact between the bacterium and host cells and this protein is also encoded by *E. faecalis* AHG0090 and other non-pathogenic isolates (e.g. *E. faecalis* PC1.1 (Ó Cuív et al., 2013), *E. faecalis* 62 (Brede et al., 2011)). Separately, the human vaginal isolate *E. faecalis* MN1 produces an NF-КB suppressive tetramic acid termed reutericyclin (Brosnahan et al., 2013). The reutericyclin regulon has been described in *Lactobacillus reuteri* and includes a non-ribosomal peptide synthetase (NRPS) and polyketide synthetase (PKS) enzymes that function in its biosynthesis (Lin et al., 2015). *E. faecalis* AHG0090 does not encode the reutericyclin regulon and nor does it encode any NRPS or PKS genes. We did not identify any Mam like sequences which is consistent with its narrow phylogenetic distribution (Quevrain et al., 2016) nor did we identify any genes that might encode the candidate <3 kDa peptidic bioactive. It is increasingly facile to produce microbial genomic sequence data but the ability to link genes with function remains challenging. It is estimated that the human gut microbiome is comprised of as much as 9.8 million non-redundant genes (Li et al., 2014). However, despite the wealth of microbial (meta)genomic data that is now publicly available the vast majority of genes remain functionally uncharacterised (Anton et al., 2013;Dantas et al., 2013). For instance, it is widely acknowledged the gut microbiota exerts a broad range of immunomodulatory activities (e.g. (Geva-Zatorsky et al., 2017)) however, with some notable exceptions (Mazmanian et al., 2005;Quevrain et al., 2016), the genes underpinning these capacities remain largely cryptic.

Microbial culturing is a time consuming and labour-intensive process although this provides the best opportunity to link genes with function. We believe focusing culturing efforts on genetically tractable strains will ultimately expedite the functional dissection of the microbiome. We previously observed the pEHR plasmids are stably maintained in their recipient hosts (Ó Cuív et al., 2015). We have now demonstrated they are maintained in *E. faecalis* AHG0090 and that this strain can be manipulated using standard molecular microbiology approaches for transformation and plasmid curing. We did identify some minor differences between the compiled pEHR vector sequences and those produced from the genome sequence data and we believe that these likely occurred during the vector construction process. We are continuing to extend the functionalities of the pEHR vector system and we anticipate that this will enable us to apply forward and/or reverse genetic approaches to functionally dissect *E. faecalis* AHG0090 and other gut microbes. For instance, NF-КB is a master regulator of inflammation and gut barrier integrity, and is central to the pathogenesis of several chronic (gut) diseases (Atreya et al., 2008;Sakamoto and Maeda, 2010;Esser et al., 2015). The gut microbiota produces a plethora of immunomodulatory bioactives and these could be used as lead molecules to catalyse the development of new biotechnologies and therapeutics. Genetic methods offer new opportunities to identify these bioactives and they complement existing -omic based methods for gene and protein function discovery (Clarke et al., 2005;Meng et al., 2012).

In conclusion, we demonstrated that metaparental mating can be used to isolate genetically tractable bacteria from the human gut that possess potent anti-inflammatory activities. Although *E. faecalis* is amongst the best characterised *Firmicutes* affiliated gut bacteria our data suggests that this taxon possesses novel anti-inflammatory capacities. Several fastidious anaerobic gut bacteria have been suggested as next generation probiotics for chronic gut diseases but *E. faecalis* may be a superior candidate due to its ease of propagation. We anticipate that the genetic dissection of *E. faecalis* AHG0090 will provide new insights into the immunomodulatory capacity of this taxon, and a deeper understanding of the early life events that help establish a tolerogenic immune response.

## Author contributions

PÓC conceived the study with MMcG, JB and MM; PÓC isolated *E. faecalis* AHG0090 and performed the genetic characterization; RG and PÓC prepared samples and performed the immunomodulatory characterizations; RG performed the immunofluorescence and gene expression experiments; ECH and PÓC performed the genome analyses; PÓC, RG, ECH, MMcG, JB and MM analyzed the data, and; PÓC wrote the manuscript with RG, ECH, MMcG, JB and MM.

## Funding

This research was supported via funds provided by the University of Faculty of Medicine (PÓC, JB and MM), the University of Queensland Diamantina Institute (MM) and the University of Queensland Microbiome Challenge Grant (MM and PÓC), as part of its contribution to the International Human Microbiome Consortium. PÓC is supported by The University of Queensland’s Reginald Ferguson Fellowship in Gastroenterology. MMcG is supported by an NHMRC Senior Research Fellowship. The Translational Research Institute is supported by a grant from the Australian Government.

## Acknowledgements

We gratefully acknowledge the Australian Centre for Ecogenomics for the library construction and genome sequencing, and Emma Hamilton-Williams for collecting the infant faecal sample.

## Conflict of interest

The authors declare that the research was conducted in the absence of any commercial or financial relationships that could be construed as a potential conflict of interest.

